# Neuronal HDAC9: A key regulator of cognitive and synaptic aging, rescuing Alzheimer’s disease-related phenotypes

**DOI:** 10.1101/2025.09.13.675847

**Authors:** Yun Lei, Yuting Chen, Ming Guo, Florikaben Patel, Yu Bai, Brandee Goo, Quansheng Du, Neal L. Weintraub, Xin-Yun Lu

## Abstract

Epigenetic regulation is a key determinant of the aging process, and its dysregulation contributes to cognitive aging and increased vulnerability to Alzheimer’s disease (AD). As major regulators of epigenetic processes, histone deacetylases (HDACs) have emerged as potential therapeutic targets for cognitive enhancement in neurodegenerative diseases. However, the distinct roles of individual HDAC isoforms remain to be defined. Here, we report that HDAC9 is specifically expressed in neurons of human and mouse brains, and its expression declines with age. HDAC9 deficiency impairs cognitive function and synaptic plasticity in young mice. Selective deletion of HDAC9 in hippocampal CA1 neurons also induces cognitive impairment. In contrast, overexpression of HDAC9 in forebrain glutamatergic neurons preserves cognitive function in aged mice. Moreover, HDAC9 is also downregulated in the brain of AD mouse models, whereas neuronal overexpression of HDAC9 alleviates AD-related cognitive and synaptic deficits and reduces Aβ deposition. Together, these findings suggest neuronal HDAC9 is necessary and sufficient for maintaining cognitive and synaptic functions in the context of aging and AD.

## INTRODUCTION

Aging is associated with cognitive decline and increased vulnerability to Alzheimer’s disease (AD), the leading cause of dementia, accounting for 60–80% of cases ^1^. While amyloid-β (Aβ) plaques and neurofibrillary tangles are the classical neuropathological hallmarks of AD, their correlation with cognitive decline is relatively modest ^2^. In contrast, synaptic dysfunction and loss are among the earliest and most consistent pathological changes in AD and show the strongest correlation with cognitive decline ^2,3^. Epigenetic modifications play a critical role in synaptic plasticity, learning, and memory formation ^4–6^. With aging, these finely tuned regulatory processes become increasingly dysregulated ^7–11^, disrupting gene expression programs that support synaptic function and cognitive performance ^11–13^. In AD, epigenomic dysfunctions further contribute to neuronal vulnerability, synaptic loss, and cognitive deficits ^12–15^.

On of the best-characterized epigenetic modifications is histone acetylation, which is dynamically regulated by histone acetyltransferases (HATs) and histone deacetylases (HDACs). While acetylation by HATs relaxes chromatin structure and facilitates transcriptional activation, HDACs promote chromatin compaction and repress gene expression ^16,17^. HDACs have been implicated in the regulation of molecular pathways underlying memory formation and synaptic plasticity ^18–20^. Pharmacological inhibition of HDACs leads to histone hyperacetylation, chromatin decondensation and transcriptional derepression ^16,17,21^. In preclinical models of AD, HDAC inhibition has been shown to enhance synaptic plasticity, improve memory, and ameliorate cognitive deficits ^22–30^. However, a major limitation of current HDAC inhibitors is their lack of isoform specificity, which can result in unintended off-target effects ^31,32^.

Mammalian HDACs are classified into four major classes: Class I (HDACs 1, 2, 3, 8), Class II (IIa: HDACs 4, 5, 7, 9; IIb: HDACs 6, 10), Class III (Sirtuins SIRT1–7), and Class IV (HDAC11) ^33,34^. Current HDAC inhibitors generally target the conserved catalytic pocket shared by zinc-dependent HDACs, resulting in limited isoform selectivity ^33,35^. Class IIa HDACs possess very weak intrinsic deacetylase activity due to the substitution of a conserved tyrosine residue essential for catalysis in other HDAC classes with a histidine residue ^36,37^. As a result, conventional HDAC inhibitors have minimal effects on Class IIa HDACs. Furthermore, no histone or other protein substrates have been definitively identified for this subclass. Instead, accumulating evidence indicates that Class IIa HDACs function primarily as transcriptional co-repressors by acting as scaffolding proteins ^33,38^. A key role of Class IIa HDACs is to recruit enzymatically active class I HDACs, such as HDAC3, to chromatin ^39–43^. Additionally, the N-terminus of Class IIa HDACs serve as a unique adaptor domain that binds diverse transcription factors, thereby modulating gene expression in a context-dependent manner ^37,44–46^. These unique structural and binding properties suggest that Class IIa HDACs exert their functions largely through deacetylase-independent mechanisms, relying on interactions with specific epigenetic modulators and transcription factors to regulate gene expression.

Given their negligible intrinsic deacetylase activity ^34^, the isoform-specific functions of Class IIa HDACs have been investigated using molecular and genetic approaches, though much remains unknown. In *Drosophila*, both overexpression and knockdown of HDAC4 impair memory, suggesting that HDAC4 is both a repressor when in excess and essential for normal memory function ^47^. In mice, HDAC4 knockout in forebrain neurons produces conflicting effects on spatial and fear memories depending on the Cre line used ^48,49^. HDAC5 knockout impairs spatial and associative memory ^50^ but does not affect memory recall or extinction ^49^. Conversely, overexpression of HDAC7 impairs long-term memory, whereas knockdown enhances it ^51^. These findings indicate the complex, context-dependent roles of Class IIa HDACs in regulating cognitive behaviors.

HDAC9, another Class IIa HDAC, is highly expressed in neurons ^52^; however, its specific function in the brain remains largely undefined. Transcriptomic analyses of postmortem brain tissues revealed a reduction in HDAC9 expression in the cortical regions of individuals with AD ^53–56^. Moreover, HDAC9 expression appears to be negatively correlated with Braak stage ^56^, indicating a potential association between lower HDAC9 levels and disease progression. In this study, we characterized HDAC9 expression in both human and mouse brains, investigated its regulation during aging and in mouse models of AD, and further examined its functional roles in cognitive performance and synaptic plasticity in young, aged, and AD mice. We show that HDAC9 is downregulated in the aging brain and in two mouse models of AD. HDAC9 deletion leads to cognitive and synaptic impairments in young mice, whereas neuron-specific overexpression preserves cognitive function in aged mice and mitigate cognitive and synaptic deficits, as well as amyloid pathology, in AD mice.

## MATERIALS AND METHODS

### Animals

Male and female wild-type (WT) C57BL/6 mice (Strain #000664), Emx1-ires-Cre mice (Strain # 005628) were purchased from Jackson Laboratory (Bar Harbor, MA, USA). Male mice used for aging-related studies were obtained from National Institute on Aging (NIA) Aged Rodent Colonies. The HDAC9 knockout mouse line on the C57BL/6 background, with the deletion of most of exon 4 and all of exon 5, was developed by Dr. Eric Olson ^44^ and maintained in our colony by backcrossing with C57BL/6J mice ^57^. Male and female HDAC9 heterozygous knockout mice (HDAC9^+/-^) were bred to generate HDAC9 homozygous knockout (HDAC9^-/-^) mice and WT littermate controls. HDAC9 floxed mice on the C57BL/6 background, in which exon 4 of the mouse HDAC9 gene is flanked by loxP sites, were generated as previously described ^58^. HDAC9 floxed mice were backcrossed with C57BL/6J mice. Breeding colonies of HDAC9^flox/flox^ mice were subsequently established to produce homozygous floxed offspring for Cre-mediated, cell- and region-specific knockout studies.

*Generation of conditional HDAC9 knock-in mice.* To generate the HDAC9 conditional overexpression mouse model, the DNA fragment encoding 3×FLAG-mouse HDAC9 (mHDAC9) was synthesized (Genewiz) and cloned into the CAG-STOP-eGFP-ROSA26TV targeting vector (CTV vector, Addgene plasmid #15912) ^59^. This vector contains a loxP-flanked stop cassette downstream of the CAG promoter, allowing cell type–specific or temporal gene expression upon Cre-mediated excision (Fig. 4A). The synthesized DNA was cloned into the unique AscI site located between the LoxP-Stop-LoxP (LSL) cassette and the Frt-IRES-eGFP-Frt-polyA cassette. The targeting construct was linearized with *AsiSI* and electroporated into G4 embryonic stem (ES) cells ^60^. Correctly targeted embryonic stem (ES) cells were identified by PCR screenings using primer sets located external to the 5’ and 3’ homologous regions (Fig. 4B). Successfully targeted clones were injected into albino C57BL/6J blastocysts, and chimeric founders were bred with C57BL/6J mice to generate germline-transmitting HDAC9-Tg^flox-STOP-flox^ heterozygotes. The PCR primers used for screening and genotyping were as follows: CTV 5’LR, forward-5’-TCCTCAGAGAGCCTCGGCTAGGT-3’, reverse-5’-CCCTGGACTACTGCGCCCTACAGATGCTAGA-3’; CTV 3’LR, forward-5’-GCATCGCATTGTCTGAGTAGGTGTCA-3’, reverse-5’-CTCGAAGACCTGTTGCTGCTCAGACAGCA-3’ and HDAC9-forward-5’-GGTGCTCTGCCTGCTTCTCAGCTGCA-3’, CTV-IRES-reverse-5’-CGCTTGAGGAGAGCCATTTGACTC-3’. Female heterozygous HDAC9-Tg^flox-STOP-flox^ mice were bred with male Emx1-ires-Cre mice to produce Emx1-ires-Cre;HDAC9-Tg^flox-STOP-flox^ mice and Emx1-ires-Cre littermate controls.

The 5xFAD and APP/PS1 transgenic mice were obtained from Mutant Mouse Resource & Research Centers (MMRRC). The 5xFAD mice overexpress mutant human amyloid beta precursor protein 695 (APP) with the Swedish (K670N, M671L), Florida (I716V), and London (V717I) familial AD mutations along with human presenilin 1 (PSEN1) harboring two familial AD mutations, M146L and L286V. Both transgenes are driven by the mouse *Thy1* promoter to direct their overexpression specifically in neurons. Male hemizygous 5xFAD mice were crossed with female heterozygous HDAC9-Tg^flox-STOP-flox^ mice to generate 5xFAD/HDAC9^flox-STOP-flox^ mice, 5xFAD mice and WT littermates. The APP/PS1 mice express a chimeric mouse/human *APP* carrying Swedish mutations (K595N/M596L) and a mutant human *PSEN1* (ΔE9) under the control of mouse prion protein promoter elements.

Mice were housed in groups of 2-5 mice per individually ventilated cage, with ad libitum access to food and water under a 12-h light-dark cycle (lights on at 06:00 AM). All animal procedures were conducted in accordance with the National Institutes of Health (NIH) Guide for the Care and Use of Laboratory Animals and approved by the Institutional Animal Care and Use Committee of Augusta University.

### Stereotaxic surgery for intra-CA1 AAV vector injection

For targeted gene deletion in the hippocampal CA1 region, AAVs were delivered via stereotaxic injection under anesthesia as previously described ^61^. AAV5-CMV-Cre-GFP (referred to as AAV-Cre-GFP) and control AAV5-CMV-GFP (referred to as AAV-GFP), each at a titer >1 × 10¹² vg/mL (UNC Vector Core, Chapel Hill, NC), were injected bilaterally into the dorsal hippocampal CA1 region (coordinates: anterior–posterior (AP) = -2.1 mm, medial–lateral (ML) = ± 1.5 mm, dorsal– ventral (DV) = −1.7 mm from the bregma) of adult male HDAC9^flox/flox^ mice using a 30-gauge Hamilton microliter syringe connected to a UMP3 UltraMicroPump (World Precision Instruments, Sarasota, FL). A total volume of 0.5 μL AAV vectors (per side) was delivered at a rate of 0.1 μL/min. After completion, the needle was left in place for 5 min to minimize backflow before being slowly withdrawn. HDAC9 floxed mice were randomly received AAV-GFP or AAV-Cre-GFP injection. Behavioral tests were conducted three weeks after AAV injection to allow sufficient time for transgene expression. At the end of the experiments, injection sites were verified in each animal by examining GFP fluorescence in brain sections. Mice with “missed” injections were excluded from statistical analyses.

### Intravenous (tail vein) injection of AAV

Mice were gently warmed to dilate the vein prior to being placed in a restraint for intravenous injection. AAV-PHP.eB.hSyn.HI.eGFP-Cre.WPRE.SV40 (AAV-PHP.eB.eGFP.Cre) was a generous gift from Dr. James M. Wilson (Addgene viral prep # 105540-PHPeB). AAV-PHP.eB-EF1a-GFP was packaged by the Viral Vector Core in the Department of Neuroscience and Regenerative Medicine at Augusta University. AAV-PHP.eB viruses (1×10^12^ vg/mL, 100 μL total volume, diluted in sterile saline) were slowly injected into the lateral tail vein using a sterile 27– 30-gauge needle attached to 0.5 mL syringe. Following injection, the needle was left in place for 3 seconds before being withdrawn, and gentle pressure was applied to the injection site to prevent bleeding.

### Behavioral procedures

#### Y-maze spontaneous alternation test

Short-term spatial working memory was assessed using a Y-maze that consisted of three arms (30 × 6 × 15 cm) positioned at 120° to each other (Panlab Harvard Apparatus, MA, USA). Spontaneous alternation behavior reflects the natural tendency of mice to explore previously unvisited arms of the maze, driven by their innate curiosity and preference for novelty. Each mouse was placed at the end of one arm, facing the center of the maze, and allowed to freely explore all three arms of the maze for 10 min. The sequence and frequency of arm entries were recorded by experimenters who were blind to genotypes and treatment. An entry was defined as all four limbs being inside an arm. The percentage of spontaneous alternation [%] was calculated using the following formula: Spontaneous Alternation (%) = [Number of consecutive entries into all three arms / (Total number of arm entries − 2)] ×100% ^62^.

#### Novel object recognition (NOR) test

The NOR test has been widely used to assess object recognition memory, which is based upon rodent’s innate tendency to explore novel objects over familiar ones ^63–65^. The test was performed in a white acrylic apparatus consisting of a 40 × 40 cm open-field arena with 40 cm high walls. This test consists of three sequential sessions: habituation session, training session and testing session. On the first day, the mouse was placed individually in the empty open arena and allowed to explore for 5 min to habituate to the testing environment (habituation session). On the second day, two identical objects were placed in the arena, located close to two adjacent corners. The mouse was released against the center of the opposite wall and allowed to freely explore for 5 min (training session). After a 2-hour retention interval, a novel object was introduced to the arena and the mouse was returned to the arena and allowed to explore the familiar and novel objects for 5 min. The time spent exploring the novel object versus the familiar one was recorded. The location of novel versus familiar object was counterbalanced between mice.

#### Social recognition memory test

The social recognition memory was evaluated using a two-session paradigm. In the social interaction session, the test mouse was placed in a rectangular arena (45 × 25 × 20 cm) containing two wire cages positioned at opposite ends: one cage contained an unfamiliar social target mouse (M1), matched in age, strain, and sex to the test mouse; the other cage was left empty. The test mouse was allowed to freely explore the arena and interact with the social target for 10 min. After a 2-hour interval, the social discrimination session was conducted. During this session, a second novel social target mouse (M2) was placed in the previously empty cage (novel mouse M1 and mouse M2 were age-matched and housed in separate cages). The test mouse was again allowed to explore the arena for 5 min. To minimize stress-related confounds, social target mice were pre-habituated to the wire cages within the testing arena for 15 min per day for three consecutive days before being used for testing. Interaction time with each social target mouse during the social discrimination session was recorded. The social recognition index (%) was calculated using the formula: Social recognition index (%) = Time_Interacting with novel mouse_ /(Time_Interacting with novel mouse_+Time_Interacting with familiar mouse_)×100%.

#### Contextual fear conditioning test

The contextual fear conditioning test is one of the most widely used paradigms to assess associative fear learning and memory ^66^. Fear conditioning was performed in a mouse fear-conditioning chamber equipped with the Coulbourn Freezeframe system. The chamber was positioned inside a sound-attenuating cabinet (Coulbourn Instruments, Whitehall, PA, USA). On Day 1, mice were placed individually into the conditioning chamber and allowed to freely explore for 3 min. This was followed by four footshocks (0.8 mA, 2 seconds each) delivered at 60-second intervals. Mice remained in the chamber for an additional 60 seconds after the final shock before being returned to their home cages and housed individually overnight. On Day 2, 24 hours after conditioning, mice were reintroduced into the same chamber for 5 min with no footshock administered. Contextual fear memory was quantified by measuring freezing behavior. Freezing time was analyzed by the Freezeframe3 software. The chamber was cleaned using 20% ethanol and dried between each mouse.

#### Hot-plate Test

To control for potential confounding effects of altered pain sensitivity on freezing behavior observed during fear conditioning, a hot-plate test was conducted to assess nociceptive response to a thermal stimulus. The hot plate was kept at a constant temperature of 55°C. Mice were individually placed on the hot plate for up to 90 seconds. The latency to exhibit a response, such as a hind paw lick or jump, was recorded. Once a response was observed, the mouse was immediately removed from the hot plate and returned to its home cage. If no response occurred, the test was terminated after 90 seconds to prevent thermal injury.

#### Locomotor activity test

Locomotor activity was measured using an open field locomotor system (Omnitech Electronics Inc, Columbus, OH) as described previously ^61^. The test apparatus consisted of a transparent acrylic box (40 × 40 × 30 cm^3^) surrounded by three sets of 16 photobeam arrays detecting horizontal (X and Y axes) as well as in the ventricle (Z axis) movements. Locomotor activity was determined by breaks in photobeams and total distance traveled was analyzed using the Fusion Software (Omnitech Electronics Inc, Columbus, OH). Mice were monitored for 30 min and the distance traveled during the first 5 min was used for statistical analysis. For 5xFAD mice, locomotor activity was analyzed by calculating the distance traveled during the habituation session in the NOR test using EthoVision XT software.

#### Elevated plus-maze test

This is a widely used anxiety test, based on the natural conflict between the drive to explore a new environment and the tendency to avoid a potentially dangerous (open) area ^67,68^. The maze was made of grey acrylic, with 4 arms (30 cm long and 5 cm wide) arranged in the shape of a “plus” sign and elevated to a height of 70 cm from the floor. Two arms have no side or end walls (open arms). The other 2 arms have side walls and end walls (12 cm high) but are open on top (closed arms). The open and closed arms intersect, having a central 5×5-cm square platform giving access to all arms. Mice were placed in the central platform facing the corner between a closed arm and an open arm, and allowed to explore the maze for 5 min. The test was recorded with a video camera mounted above the maze and connected to a computer. The time spent on the open and closed arms and the number of entries made into each arm were scored. Entry was defined as all four paws entering one arm. The degree of anxiety was assessed by calculating the percentage of open arm time (time spent in the open arms/total time spent in all arms).

### Immunohistochemistry

Mice were transcardially perfused using 0.1 M phosphate-buffered saline (PBS) followed by 4% paraformaldehyde (PFA) in PBS and processed as described previously ^69^. Briefly, 40-µm coronal sections were rinsed with 0.05 M Tris-buffered saline (TBS) for 10 min and incubated in TBS containing 0.5% (v/v) Triton-X 100 (TBST) for 1 h. Brain sections were further incubated in blocking buffer (10% donkey serum in TBST) for 1.5 h at room temperature and then incubated with rabbit monoclonal anti-HDAC9 antibody (Abcam, #ab109446, 1:400) in combination with one of cell-type specific antibodies (mouse monoclonal anti-NeuN antibody, MilliporeSigma, #MAB377, 1:500; goat polyclonal anti-IBA1antibody, FUJIFILM Wako Pure Chemical Corporation, #011-27991, 1:250; mouse monoclonal anti-GFAP antibody, cell signaling technology, #3670, 1:250) for double labeling. For Aβ immunostaining, rabbit monoclonal anti-Aβ 6E10 antibody was used (Novus Biologicals, #NBP2-62566, 1:750). These primary antibodies were incubated with brain sections in blocking buffer at 4 °C overnight. Brain sections were rinsed in TBS for four times (10 min each) and then incubated with Alexa Fluor conjugated secondary antibodies (Alexa Fluor 488 conjugated donkey anti-rabbit, #A-21206; Alexa Fluor 647 conjugated donkey anti-mouse, # A-31571; Alexa Fluor 555 conjugated donkey anti-goat, # A-21432; Alexa Fluor 555 conjugated donkey anti-mouse, # A-31570; 1:500) at room temperature for 2 h. The sections were further stained with DAPI, washed with TBS and then mounted on slides with aqueous anti-fade mounting medium. KEYENCE All-in-One Fluorescence Microscope BZ-X800 (Keyence Corporation of America, Itasca, IL) and Nikon A1R MP+ Multiphoton/Confocal Microscope were used to visualize immunostaining and capture the images. Quantitative analysis of Aβ plaque load was conducted using Fiji software as previously described ^70,71^. Briefly, a consistent intensity threshold was set across samples to identify specific staining within the selected brain regions, and Aβ plaque load was defined as the percentage of the area covered by Aβ plaques (% area).

Frozen human brain tissue was transferred from -80 °C to -20°C in a cryostat to allow gradual temperature equilibration and then completely embedded in Epredia M-1 embedding matrix. Embedded tissue blocks were then cryosectioned at a thickness of 10 µm and the sections were mounted onto gelatin-coated histological slides. The tissue sections were fixed in 4% PFA at room temperature for 15 min, followed by three washes in TBS (10 min each). The area surrounding the tissue was outlined using a hydrophobic barrier pen to confine reagents during staining. Sections were permeabilized in TBST for 1 h at room temperature, followed by blocking in TBST containing 10% donkey serum for another hour. Then brain sections were incubated with rabbit monoclonal anti-HDAC9 antibody (Abcam, #ab109446, 1:200) in combination with one of following antibodies: mouse monoclonal anti-NeuN antibody (MilliporeSigma, #MAB377, 1:200), goat polyclonal anti-IBA1antibody (Abcam, #ab5076, 1:200) or mouse monoclonal anti-GFAP antibody (cell signaling technology, #3670, 1:200) diluted in TBST containing 10% donkey serum overnight at 4 °C. The following day, sections were washed four times with TBS (10 min each), then incubated for 2 h at room temperature with Alexa Fluor-conjugated secondary antibodies diluted in TBST (Alexa Fluor 488 donkey anti-rabbit, Thermo Fisher, #A-21206; Alexa Fluor 647 donkey anti-mouse, #A-31571; Alexa Fluor 555 donkey anti-goat, #A-21432; 1:500). After a brief rinse in TBS, nuclei were counterstained with DAPI, followed by three washes in TBS (10 min each). The residual buffer was carefully removed from slides and anti-fade mounting medium was applied before placing coverslips. Fluorescence imaging was performed as described above.

### RNA extraction and real-time quantitative PCR analysis

Mice were sacrificed by rapid decapitation and the hippocampus and prefrontal cortex were quickly dissected on ice. Tissues were snap-frozen in liquid nitrogen and stored in -80 °C until further processing for RNA extraction. Total RNA was extracted with the RNeasy Plus Mini Kit (Qiagen #74134, Germantown, MD). RNA purity and concentration of purified RNA were determined using a NanoDrop™ 8000 Spectrophotometer (Thermo Fisher Scientific, Waltham, MA). cDNA was synthesized from total RNA using the High-Capacity cDNA Reverse Transcription Kit (Thermo Fisher Scientific, #4368814) with random primers. The cDNA product was diluted and processed for real-time PCR quantification using the QuantStudio 5 real-time PCR system (Thermo Fisher Scientific, Waltham, MA). Gene expression levels were quantified using the following primers: *HDAC9*, forward-5′-GCGAGACACAGATGCTCAGAC-3′, reverse-5′-TGGGTTTTCCTTCCATTGCT-3′; *Bdnf* exon-IX, forward-5′-GCGCCCATGAAAGAAGTAAA-3′, reverse-5′-TCGTCAGACCTCTCGAACCT-3′; *Bdnf*exon-IV, forward-5′-CAGAGCAGCTGCCTTGATGTT-3′, reverse-5′-GCCTTGTCCGTGGACGTTTA-3′; *Bdnf* exon-VI, forward-5′-CTGGGAGGCTTTGATGAGAC-3′, reverse-5′-GCCTTCATGCAACCGAAGTA-3′; *β-actin*, forward-5′-GATCATTGCTCCTCCTGAGC-3′, reverse-5′-ACTCCTGCTTGCTGATCCAC-3′; *β-tubulin*, forward-5′-AGCAACATGAATGACCTGGTG-3′, reverse-5′-GCTTTCCCTAACCTGCTTGG-3′. As described previously ^61,69^, the 2^(-ΔΔCT)^ method was used to obtain relative fold-change of gene expression normalized by housekeeping genes.

### Western blot

Prefrontal cortex and hippocampus were dissected on ice, snap-frozen in liquid nitrogen and stored at -80 °C before being processed for western blotting. Tissues were sonicated with lysis buffer (75 mM Tris, 15% v/v glycerol, 3% w/v SDS, PH 6.8) containing protease inhibitors. Samples were lysed on rotator at 4 °C for 30 min and then centrifuged at 14, 000 rpm for 15 min at 4 °C. The protein concentration of supernatant was determined using BCA assay. Protein sample was mixed with 6×loading buffer and incubated at 70 °C for 10 min. Denatured proteins were separated on 10% Precast Protein Gels and transferred to activated polyvinylidene fluoride membrane. After transferring, the membrane was incubated with Intercept (TBS) Blocking Buffer for 1 hour at room temperature and then incubated with primary antibodies overnight at 4 °C. Anti-HDAC9 (Cat. #orb214926, Biorbyt, 1:500), mouse monoclonal anti-FLAG antibody (Cat. #F1804, Millipore Sigma, 1:1000) and anti-β-actin antibody (Cat. #A5441, Millipore Sigma, 1:5000) were diluted by Li-Cor Intercept T20 (TBS) Antibody Diluent. After washing, membrane was incubated with IRDye 800CW Donkey anti-Rabbit Secondary Antibody (1:10,000) or IRDye® 680RD Donkey anti-Mouse IgG Secondary Antibody (1:10,000) at room temperature for 1 hour. After washing three times (10 min each), signals were visualized with Odyssey CLx Imaging System (LI-COR Biosciences) and quantitatively analyzed with Image J.

### Synaptic plasticity/long-term potentiation in the hippocampus

Mice were anesthetized with isoflurane and brains were quickly transferred to an ice-cold solution (254 mM sucrose, 3 mM KCl, 2 mM MgCl_2_, 2 mM CaCl_2_, 1.25 mM NaH_2_PO_4_, 10 mM D-glucose, and 24 mM NaHCO_3_). To prepare hippocampal slices, horizontal sections (400 μm thick) at the middle septotemporal levels were cut in ice-cold choline solution using a Leica VT1000S vibratome (Leica Microsystems) and allowed to recover at 32 °C for at least 1 h in an oxygenated (95% O_2_/5% CO_2_) artificial cerebrospinal fluid (aCSF) solution (124 mM NaCl, 2 mM KCl, 2 mM MgSO_4_, 2 mM CaCl_2_, 1.25 mM NaH_2_PO_4_, 26 mM NaHCO_3_, and 10 mM Glucose, with pH 7.3 and osmolarity 300 mOsm/L). Brain slices were transferred to an interface chamber and superfused with aCSF saturated with 95% O_2_/5% CO_2_. Field excitatory postsynaptic potentials (fEPSPs) were recorded with glass electrodes in the stratum radiatum layer of CA1, in response to stimulation of the Schaffer collaterals using a bipolar concentric electrode. fEPSPs were recorded and filtered (low pass at 2 kHz) with a MultiClamp 700B amplifier (Axon Instruments, Union City, CA, USA), digitized at 10 kHz with an A/D converter (Digidata 1550A, Axon Instruments), and analyzed with Pclamp10.7 and Clampfit10.7 software (Molecular Devices, San Jose, CA, USA). Input/output for fEPSPs were measured by applying a series of stimuli with increasing intensity to the Schaffer collaterals (0.1-1.0 mA; 0.1 ms stimulation with 30 second interval). The stimulation intensity evoking 30∼50% of the maximal fEPSP slope was utilized for LTP induction. Baseline fEPSPs were recorded for a minimum of 10 min. LTP was induced by high-frequency stimulation (HFS) of the Schaffer collaterals using a single 1-sec train at100 Hz and recorded for 60 min. Data was analyzed by measuring the slope of individual fEPSPs.

### Statistical analysis

All results are presented as mean ± standard error of the mean (SEM). Statistical analyses were performed using the GraphPad Prism 10.0 software (GraphPad Software, Inc., La Jolla, CA). The Shapiro–Wilk test and the F test were utilized to assess the assumptions of normality and equal variance, respectively. For comparisons between two groups with normally distributed data and equal variances, two-tailed unpaired *t*-tests were performed. For comparisons among three groups, one-way analysis of variance (ANOVA) followed by Bonferroni post hoc test was used. For non-normally distributed data, the Mann–Whitney *U* test was used for two-group comparisons, and the Kruskal–Wallis test followed by Dunn’s multiple comparisons was applied for comparisons across multiple groups. In the NOR test and the social recognition test, exploration time of the novel versus familiar object or mouse was analyzed using a two-tailed paired *t*-test for normally distributed data or the Wilcoxon matched-pairs signed-rank test for non-normally distributed data. A *P*-value < 0.05 was considered statistically significant for all analyses.

## RESULTS

### HDAC9 is expressed in neurons and downregulated with aging in the PFC and hippocampus

The medial prefrontal cortex (mPFC) and hippocampus play an important regulatory role in cognitive functions, and both regions are particularly vulnerable to neurodegeneration in normal aging and AD ^72–74^. The rodent mPFC is functionally homologous to the dorsolateral PFC (DLPFC) in humans ^75^. To determine cell-type specific expression of HDAC9 in the brain, we performed immunostaining with a specific antibody against HDAC9 using sections of the DLPFC (Brodmann’s area 9) from human post-mortem brains. HDAC9 colocalized with the neuronal marker NeuN, but not with the astrocyte marker GFAP or the microglia marker IBA1 (Fig. 1A), suggesting HDAC9 is selectively expressed in neurons. In the mouse brains, we examined HDAC9 expression in the mPFC and hippocampus (Fig. 1B). Consistent with the observations in human brain tissue, HDAC9 was expressed in neurons but was not detected in astrocytes or microglia in either brain region (Fig. 1B).

**Figure 1.**
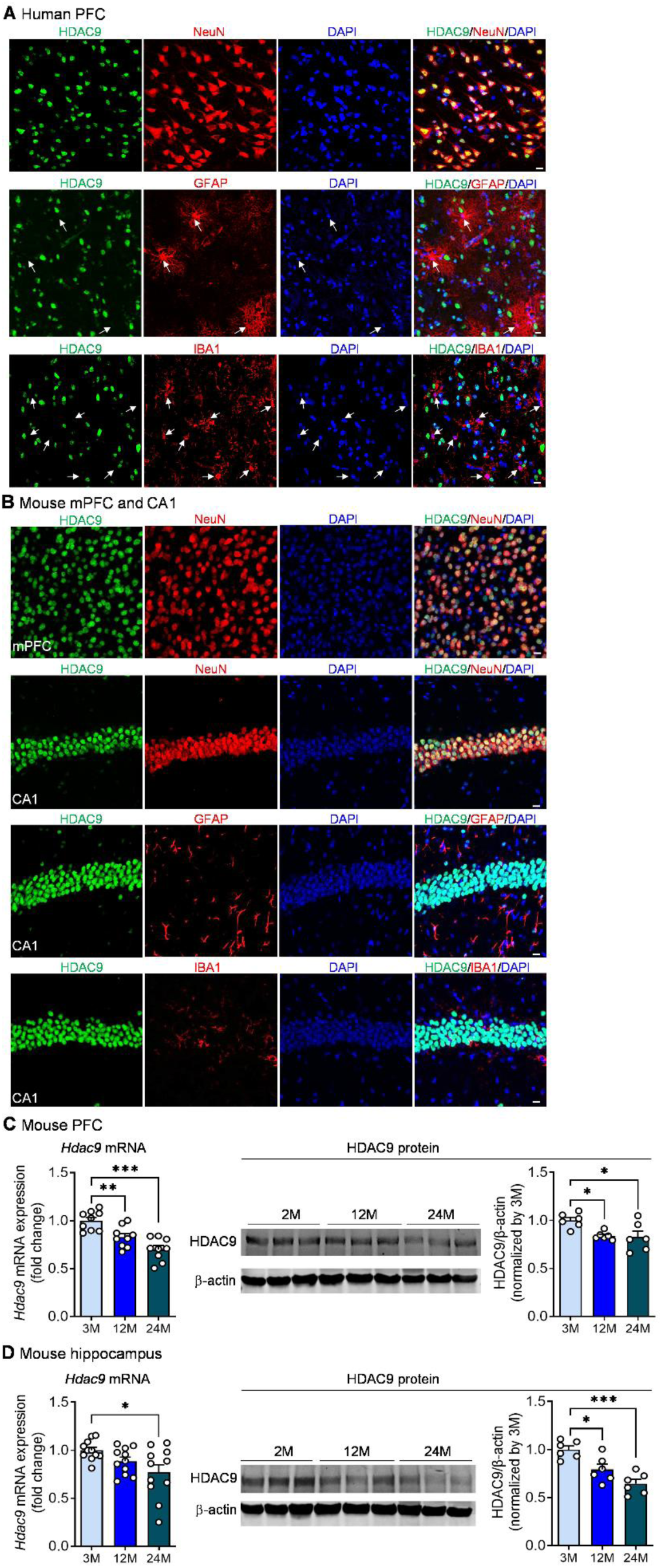
HDAC9 expression in the brain and its regulation by aging. (**A**) Double immunostaining with antibodies against HDAC9 and cell-type-specific markers: NeuN (neuronal marker), GFAP (astrocyte marker), and IBA1 (microglia marker) in the dorsolateral prefrontal cortex (Brodmann area 9) of the human brain. White arrows indicate astrocytes or microglia lacking HDAC9 expression. Scale bar = 10 μm. (**B**) Double immunostaining with antibodies against HDAC9 and NeuN, GFAP or IBA1 in the medial prefrontal cortex (mPFC) and hippocampal CA1 region of the mouse brain. Scale bar = 10 μm. (**C**) RT-qPCR and western blot detection of HDAC9 expression in the PFC of mice at different ages. mRNA expression: one-way ANOVA, *F*_(2, 24)_ = 16.670, *P* < 0.001, *n* = 9 male mice per group. Protein expression: one-way ANOVA, *F*_(2, 15)_ = 5.565, *P* = 0.016, *n* = 6 male mice per group. (**D**) RT-qPCR and western blot detection of HDAC9 expression in the hippocampus of mice at different ages. mRNA expression: Brown-Forsythe ANOVA, *P* = 0.027, *n* = 11 male mice per group. Protein expression: one-way ANOVA, *F*_(2, 15)_ = 14.860, *P* < 0.001, *n* = 6 male mice per group. \**P* < 0.05, \*\**P* < 0.01, \*\*\**P* < 0.001.

We next investigated the effects of aging on HDAC9 expressions in these two brain regions. The PFC and hippocampus were collected from wild-type mice at three different ages: 3, 12, and 24 months. Quantitative real-time PCR analysis revealed a significant age-dependent downregulation of HDAC9 mRNA expression in both the PFC and hippocampus (Fig. 1C-D). Consistently, western blotting showed a corresponding decrease in HDAC9 protein levels in these two brain regions with advancing age (Fig. 1C-D).

### HDAC9 deficiency causes cognitive deficits in young mice

To investigate whether age-dependent downregulation of HDAC9 contributes to cognitive decline, we assessed the cognitive performance of global HDAC9 knockout (HDAC9^-/-^) mice and their wild-type (WT) littermates at young ages (4-6 months old). Short-term spatial working memory was measured using the Y-maze spontaneous alternation test. Spontaneous alternation, driven by an innate curiosity to explore previously unvisited arms, serves as an indicator of spatial working memory. HDAC9^-/-^ mice exhibited significantly lower percentage of spontaneous alternation compared to WT mice, without any change in total arm entries (Fig. 2A), indicating impaired spatial working memory. To evaluate object recognition memory, we used the novel object recognition test to assess the ability to recall and differentiate between familiar and novel objects. Mice were tested for short-term object recognition memory after a 2-hour retention interval following the training phase. While WT mice spent significantly more time exploring the novel object than the familiar one, HDAC9⁻/⁻ mice showed no preference (Fig. 2B). The difference in exploration time between the novel and familiar objects was normalized to total investigation time and expressed as a discrimination index, which was significantly lower in HDAC9^⁻/⁻^ mice compared to WT controls (Fig. 2B), indicating impaired object recognition memory.

**Figure 2.**
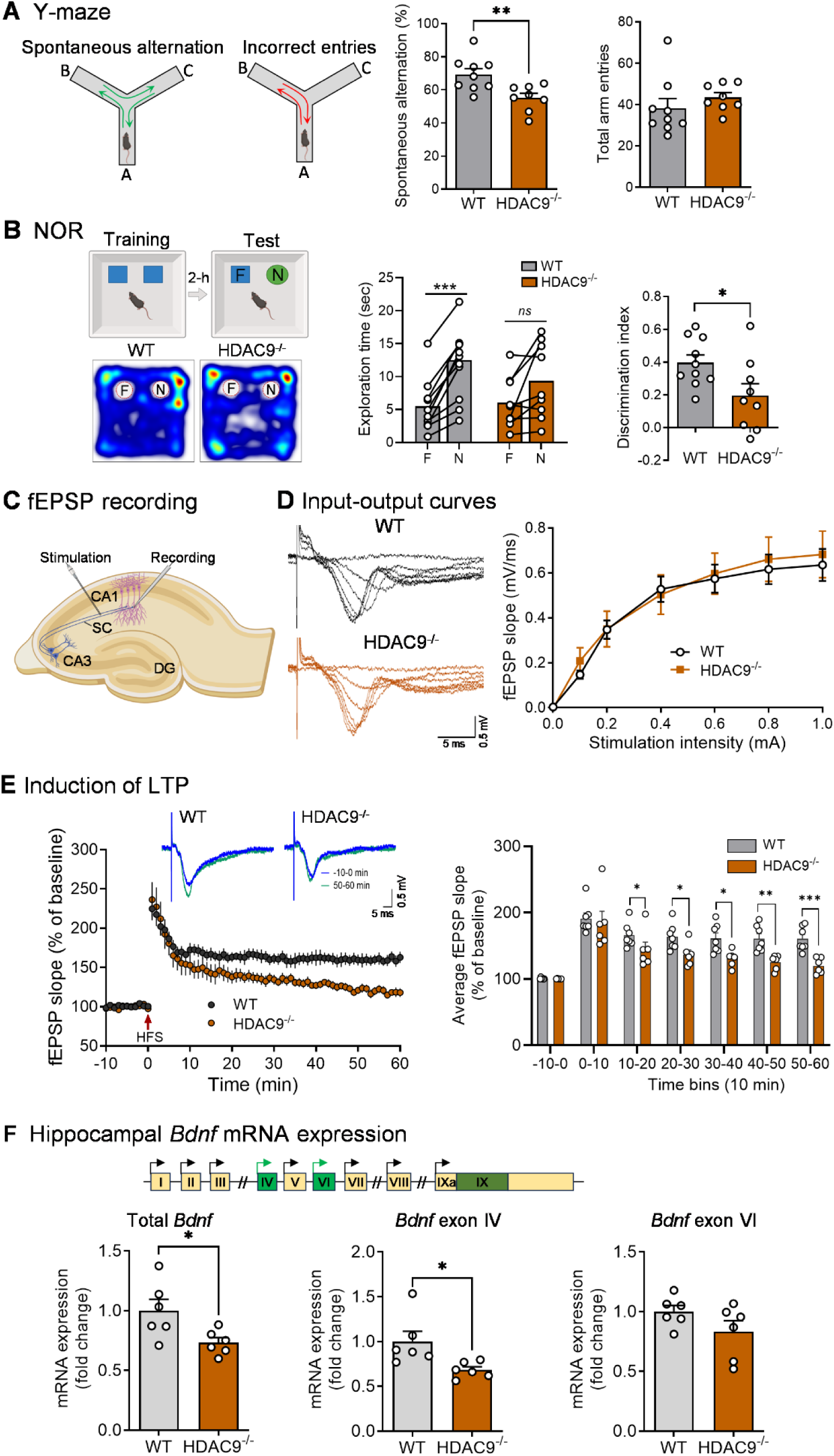
Cognitive and synaptic impairments in young HDAC9 knockout mice. HDAC9^-/-^mice and wild-type (WT) littermate controls aged 4–6 months were subjected to a battery of behavioral tests to assess cognitive function. (**A**) Y-maze spontaneous alternation test. Left, schematic of the Y-maze. Middle, spontaneous alternation rate (two-tailed unpaired t test, *t*_15_ = 3.159, *P* = 0.007). Right, total arm entries (two-tailed unpaired t test, *t*_15_ = 0.971, *P* = 0.347). WT: *n* = 9 (4 males and 5 females); HDAC9^-/-^: *n* = 8 (3 males and 5 females). (**B**) Novel object recognition (NOR) test. Left, schematic of the NOR experimental procedure (top) and representative heat maps showing time spent in the test arena (bottom). Middle, time spent exploring the familiar object (F) and the novel object (N). WT: two-tailed paired t-test, *t*_9_ = 7.150, *P* < 0.001; HDAC9^-/-^: two-sided paired t-test, *t*_8_ = 2.299, *P* = 0.051. Right, discrimination index. Two-tailed unpaired t test, *t*_17_ = 2.393, *P* = 0.029. WT: *n* = 10 (6 males and 4 females); HDAC9^-/-^: n=9 (5 males and 4 females). (**C**) Recording of fEPSPs in the CA1 region of hippocampal slices. A schematic of a transverse hippocampal slice illustrates stimulation of the Schaffer collateral pathway (SC) and placement of the recording electrode in the stratum radiatum of CA1 to obtain extracellular responses. DG, dentate gyrus. Illustrations were created with BioRender.com. (**D**) Input–output curves of fEPSPs. Left, representative traces of fEPSPs in response to increasing stimulation intensities. Right, quantitative input–output curves showing the relationship between stimulation intensity and fEPSP slope. Two-way repeated measures ANOVA: genotype, *F*_(1, 14)_ = 0.053, *P* = 0.821; stimulation, *F*_(6, 84)_ = 94.23, *P* < 0.001; genotype × stimulation, *F*_(6, 84)_ = 0.365, *P* = 0.899. (**E**) Long-term potentiation (LTP) at CA3–CA1 synapses. Left, time course of fEPSPs slopes recorded before and after high-frequency stimulation (HFS) of Schaffer collaterals. Inset, representative traces recorded 10 min before HFS (baseline) and 50 min after HFS. Two-way repeated measures ANOVA: genotype, *F*_(1, 11)_ = 6.933, *P* = 0.023; time, *F*_(70, 770)_ = 42.65, *P* < 0.001; genotype × time, *F*_(70, 770)_ = 3.533, *P* < 0.001. Right, average fEPSP slope calculated in 10-min time bins before and after LTP induction. Mann Whitney test: -10-0 min, *P* = 0.731; 0-10 min, *P* = 0.445; 10-20 min, *P* = 0.035. Two-tailed unpaired t test: 20-30 min, *t*_11_ = 2.593, *P* = 0.025; 30-40 min, *t*_11_ = 3.061, *P* = 0.011; 40-50 min, *t*_11_ = 3.894, *P* = 0.003; 50-60 min, *t*_11_ = 4.627, *P* < 0.001. WT: *n* = 7 slices from 4 mice (2 males and 2 females); HDAC9^-/-^: *n* = 6 slices from 4 mice (2 males and 2 females). (**F**) *Bdnf* expression in the hippocampus. Top, structure of the mouse *Bdnf* gene. Bottom, expression levels of total *Bdnf* (two-tailed unpaired t test, t_10_ = 2.547, P = 0.029), *Bdnf* exon-IV (unpaired t test with Welch’s correction, P = 0.038) and *Bdnf* exon-VI transcripts (two-tailed unpaired t test, t_10_ = 1.587, P = 0.144). *n* = 6 mice per group. \**P* < 0.05, \*\**P* < 0.01, \*\*\**P* < 0.001.

### HDAC9 deficiency impairs hippocampal synaptic plasticity

Long-term synaptic potentiation (LTP), a form of synaptic plasticity, is widely regarded as one of the primary cellular mechanisms underlying learning and memory, particularly in the hippocampus, a brain region critical for encoding and consolidating new memories ^76,77^. Notably, impaired LTP is strongly associated with memory deficits observed in a range of pathological and physiological conditions, including aging and AD ^78^. LTP is most commonly recorded in the CA1 region of the hippocampus due to its well-defined monosynaptic pathway from CA3 pyramidal neurons via the Schaffer collaterals, which allows for precise stimulation and reliable recording ^79^. HDAC9 is abundant throughout the hippocampus including the CA1 region (Fig. 1B). To determine whether HDAC9 is necessary for synaptic plasticity in the hippocampus, extracellular recordings of field excitatory postsynaptic potentials (fEPSPs) were conducted in the CA1 area of hippocampal slices from HDAC9^⁻/⁻^ mice and their WT littermate controls, specifically targeting synapses in the stratum radiatum, where Schaffer collateral inputs from CA3 neurons terminate on the apical dendrites of CA1 pyramidal neurons (Fig. 2C) ^79^. First, basal synaptic transmission was assessed by generating input-output relationships by plotting the slope of the fEPSP in response to a series of increasing presynaptic stimulation intensities. HDAC9^-/-^ mice exhibited input-output curves that were comparable to those of WT controls (Fig. 2D), indicating that basal synaptic transmission remains intact in the absence of HDAC9. The slope of the fEPSP increased proportionally with presynaptic stimulation intensity in both genotypes. These results demonstrate that deletion of HDAC9 does not disrupt the basic input-output relationship between presynaptic input and postsynaptic response. Next, we examined LTP in hippocampal slices from HDAC9^-/-^ mice and WT controls to evaluate activity-dependent synaptic plasticity. LTP was induced at CA3–CA1 hippocampal synapses by high-frequency stimulation of the Schaffer collaterals using a train (1s duration) of 100 Hz pulses, and fEPSPs were recorded in the stratum radiatum of the CA1 region. Immediately following high-frequency stimulation, HDAC9^⁻/⁻^ mice and WT controls exhibited comparable initial synaptic potentiation, indicating that LTP induction was intact in both groups (Fig. 2E). This was followed by a sharp decline in potentiation within the first 10 min, consistent with the expected early decay phase of LTP. After this initial drop, WT mice maintained a stable level of potentiation throughout the recording period, whereas HDAC9^⁻/⁻^ mice continued to decline, with synaptic responses approaching baseline levels by the end of the session (Fig. 2E). A significantly lower magnitude of LTP was evident in HDAC9^⁻/⁻^ mice after 10 min, and the difference between the two groups became progressively larger over time (Fig. 2E). These findings suggest that HDAC9 is not required for the maintenance of fundamental synaptic strength and the induction of LTP, but it is critically involved in the stabilization and maintenance of synaptic plasticity. Thus, HDAC9 likely plays a key role in sustaining activity-dependent synaptic modifications necessary for memory storage and persistent circuit remodeling.

Brain-derived neurotrophic factor (BDNF) is a key modulator of synaptic plasticity and memory formation, and its dysregulation has been implicated in both aging-related cognitive decline and AD ^80–82^. The rodent *Bdnf* gene consists of nine distinct 5′ non-coding exons (I–IXa), each driven by its own promoter and spliced to a common 3′ coding exon (IX). Although all transcript variants encode the same BDNF protein, their initiation from unique upstream promoters enables precise spatiotemporal and activity-dependent regulation of *Bdnf* expression ^83^. Notably, the translation of *Bdnf* transcripts containing *Bdnf* exon-IV and -VI is regulated by neuronal activity ^84,85^ and the dysregulation of these translation is linked memory loss ^80,86^. To determine whether HDAC9 deficiency influences BDNF expression, we measured total *Bdnf* mRNA levels in the hippocampus of HDAC9^⁻/⁻^ mice and their WT littermate controls at 4-6 months of age. As shown in Fig. 2F, total *Bdnf* mRNA levels were significantly reduced in HDAC9^⁻/⁻^ mice. This reduction may reflect changes in the expression of specific transcript variants arising from non-coding exons. Indeed, in the hippocampus of HDAC9^⁻/⁻^ mice, Bdnf exon-IV transcript levels were significantly decreased, whereas exon-VI transcript levels remained unchanged compared to WT controls (Fig. 2F).

### CA1-specific HDAC9 deletion impairs cognitive function in young mice

To determine the role of HDAC9 specifically within the hippocampal CA1 region, we employed the Cre/loxP system to induce region-specific deletion of HDAC9 in adult mice (4-6 months old). Adult HDAC9^flox/flox^ mice (Fig. 3A) received bilateral microinjections of AAV-Cre-GFP or AAV-GFP into the CA1 region (Fig. 3B). Cre recombinase and GFP expression in these vectors were driven by the human synapsin (hSyn) promoter, conferring neuron-specific expression. Behavioral tests were performed 3 weeks after AAV injection. In the Y-maze, mice that were injected with AAV-Cre-GFP exhibited impaired spatial working memory, as evidenced by a reduced spontaneous alternation rate in the Y-maze test compared to control mice injected with AAV-GFP (Fig. 3C). Additionally, AAV-Cre-GFP-injected mice showed increased total arm entries, suggesting hyperactivity induced by CA1-specific HDAC9 deletion. In the novel object recognition test, control mice (AAV-GFP) displayed a significant preference for the novel object, whereas AAV-Cre-GFP-injected mice failed to discriminate between novel and familiar objects. The discrimination index was reduced in AAV-Cre-GFP-injected mice (Fig. 3D), indicating a deficit in recognition memory associated with CA1-specific HDAC9 loss.

**Figure 3.**
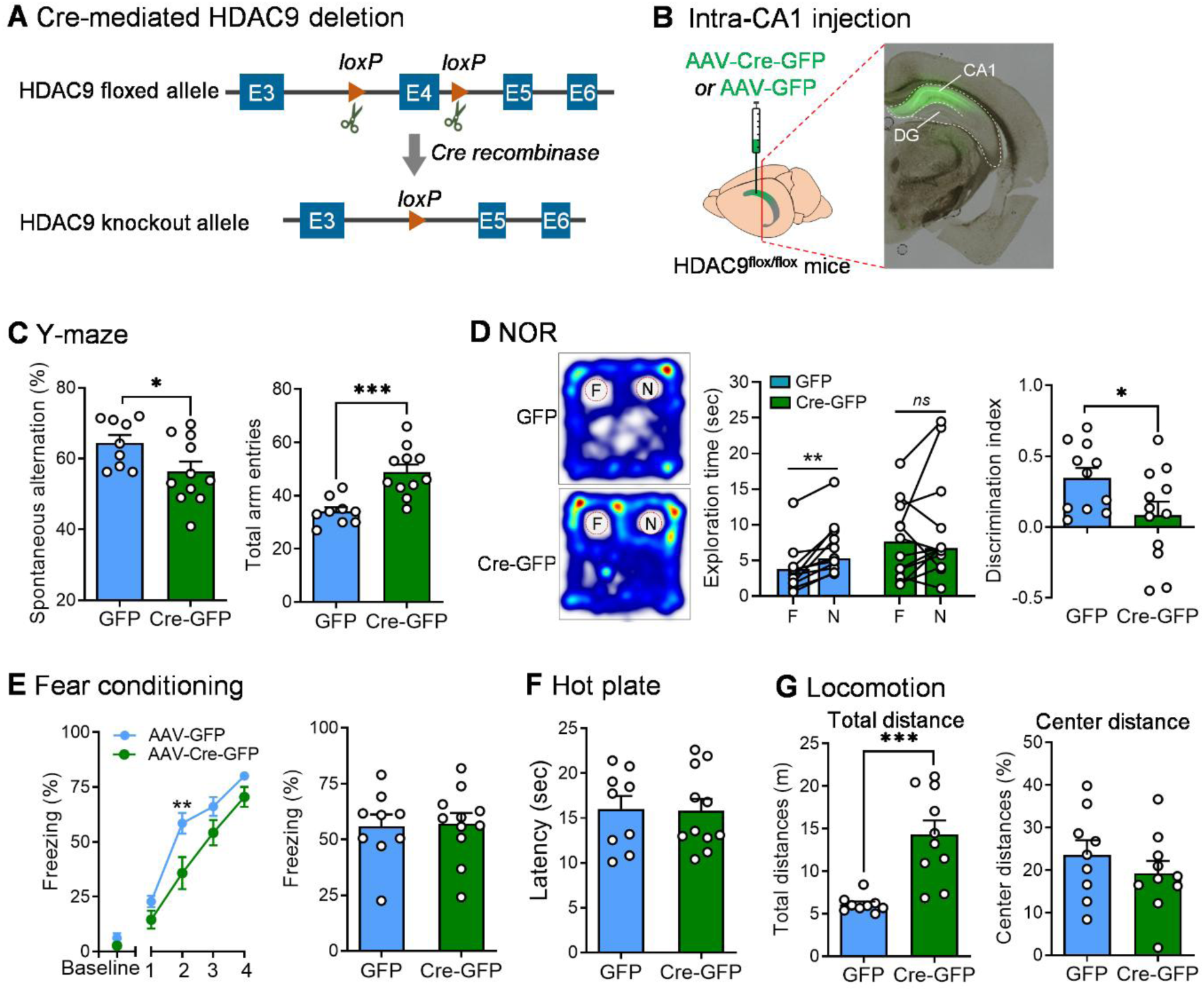
Cognitive deficits in adult HDAC9^flox/flox^ mice with CA1-specific deletion of HDAC9 at 4-6 months of age. (**A**) Schematic of conditional HDAC9 deletion mediated by Cre recombinase in HDAC9^flox/flox^ mice. (**B**) Schematic illustration of stereotaxic injection of AAV-Cre-GFP or AAV-GFP vectors into the CA1 region and representative image showing GFP expression at the injection site in CA1. (**C**) Y-maze spontaneous alternation test. Left, spontaneous alternation rate: two-tailed unpaired t test, *t*_18_ = 2.178, *P* = 0.043. Right, total arm entries: two-tailed unpaired t test, *t*_18_ = 4.370, *P* < 0.001. AAV-GFP: *n* = 9 mice; AAV-Cre-GFP: *n* = 11 mice. (**D**) Novel object recognition (NOR) test. Left, representative heat maps showing time spent in the test arena. Middle, time spent exploring the familiar object (F) and the novel object (N): AAV-GFP: Wilcoxon matched-pairs signed-rank test, *P* = 0.001; AAV-Cre-GFP: Wilcoxon matched-pairs signed-rank test, *P* = 0.267. Right, discrimination index: two-tailed unpaired t test, *t*_21_ = 2.185, *P* = 0.040. AAV-GFP: *n* = 11 mice; AAV-Cre-GFP: *n* = 12 mice. (**E**) Fear conditioning test. Left, percentage of freezing during the fear acquisition phase. Two-way repeated measures ANOVA: treatment, *F*_(1, 18)_ = 5.799, *P* = 0.027; trial, *F*_(4, 72)_ = 148.9, *P* < 0.001; treatment × trial, *F*_(4, 72)_ = 2.204, *P* = 0.077. Right, percentage of freezing during the retrieval phase, indicating contextual fear memory 24 hours after conditioning. Two-tailed unpaired t-test, *t*_18_ = 0.163, *P* = 0.873. (**F**) Latency of paw withdrawal in the hot plate test, assessing pain sensitivity. Two-tailed unpaired t-test, *t*_18_ = 0.11, *P* = 0.912. AAV-GFP: *n* = 9 mice; AAV-Cre-GFP: *n* = 11 mice. (**G**) Locomotor activity. Left, total distance traveled in the testing arena. Unpaired t-test with Welch’s correction, *P* < 0.001. Right, percentage of distance traveled in the center of the arena. Two-tailed unpaired t-test, *t*_17_ = 0.977, *P* = 0.342. AAV-GFP: *n* = 9 mice; AAV-Cre-GFP: *n* = 10 mice. \**P* < 0.05, \*\**P* < 0.01, \*\*\**P* < 0.001, *ns*: no statistical significance.

To evaluate associative learning and memory, we conducted contextual fear conditioning, a hippocampus-dependent task. Mice received 4 electric foot shocks in the chamber and were returned to the context 24 hours after training to test for long-term memory of the context-shock association by measuring their freezing behavior (a common fear response). AAV-Cre-GFP-injected mice exhibited less freezing behavior compared to controls during the acquisition phase, but they displayed normal fear memory 24 hours after conditioning (Fig. 3E). The reduced freezing response during acquisition could indicate weak associative learning or a diminished ability to process sensory stimuli. To assess whether the observed reduction in freezing response was due to altered pain sensitivity, mice were subjected to the hot plate test. No significant differences in paw withdrawal latency were detected between groups (Fig. 3F), suggesting that the decreased freezing during conditioning reflects impaired learning, not altered sensory sensitivity. Consistent with the increased arm entries observed in the Y-maze test, AAV-Cre-GFP-injected mice displayed heightened locomotor activity, as evidenced by a greater total distance traveled in the open field test (Fig. 3G). However, no differences were observed in the percentage of distance traveled in the center zone, a common measure of anxiety-like behavior, indicating that HDAC9 deletion in CA1 does not affect anxiety levels (Fig. 3G).

### HDAC9 overexpression in forebrain glutamatergic neurons preserves cognitive function in aged mice

Given that HDAC9 expression decreases with aging in the PFC and hippocampus, and HDAC9 knockout results in cognitive impairments, we sought to determine whether HDAC9 overexpression could ameliorate age-related cognitive decline. However, the large size of the HDAC9 expression cassette precludes its delivery via AAV vectors. Therefore, we generated a conditional HDAC9 knock-in mouse model. In this model, the mouse HDAC9 (mHDAC9) cDNA, tagged at the N-terminus with 3×FLAG, was placed under the control of the CAG promoter and preceded by a loxP-STOP-loxP cassette. This design prevents transgene expression until Cre recombinase excises the STOP sequence, thereby enabling conditional, tissue-specific, or temporally regulated expression of HDAC9 (Fig. 4A). The construct was targeted to the Rosa26 locus, a “safe harbor” locus for gene targeting in mouse models due to its ubiquitous transcriptional activity, lack of protein-coding function, and minimal risk of disrupting endogenous genes, thereby avoiding side effects on normal cellular and tissue functions ^87^ (Fig. 4A). Targeting this locus enables locus-specific, copy number-controlled, and consistent transgene expression without disrupting the structure or function of endogenous genes.

**Figure 4.**
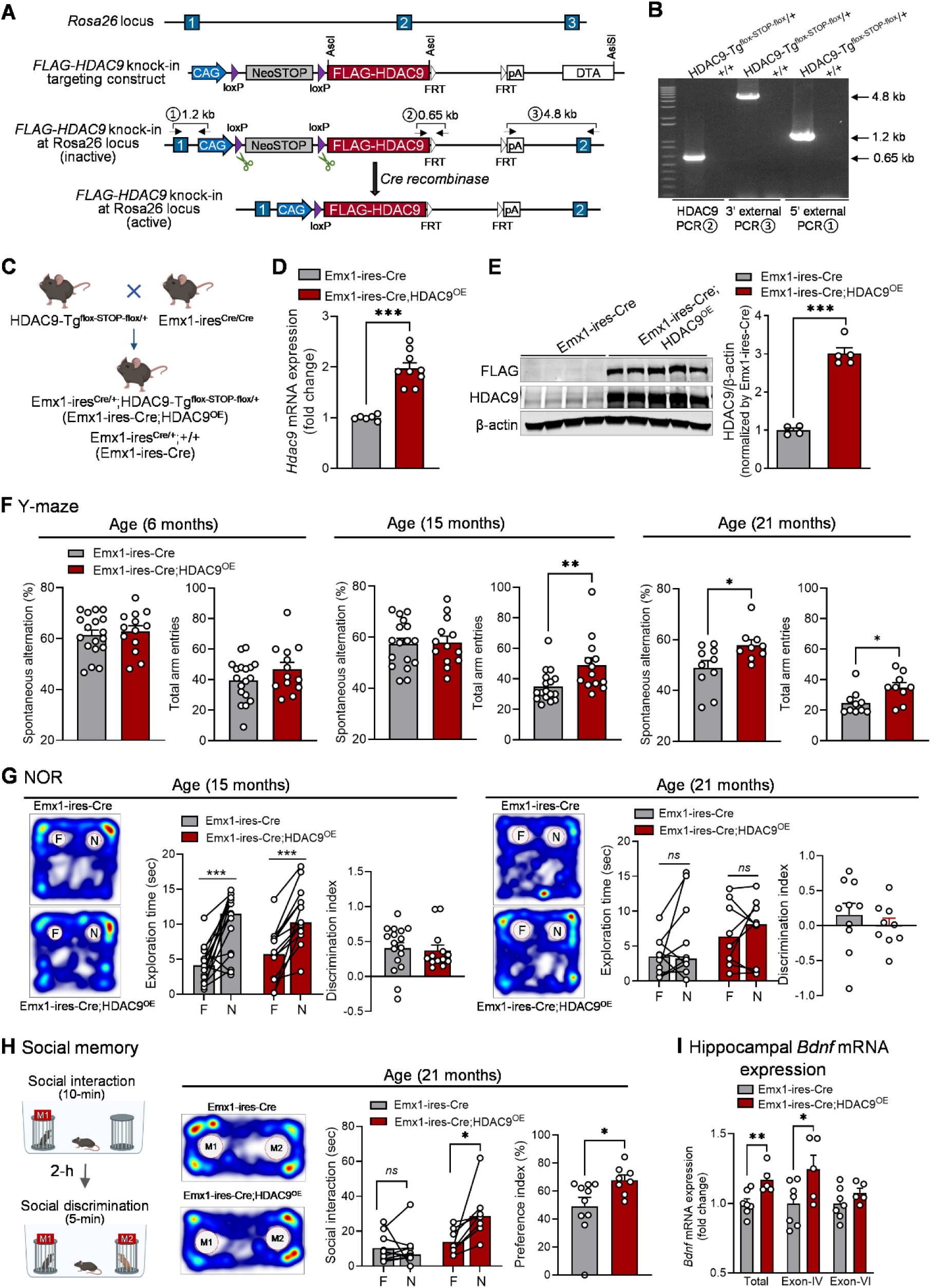
Preservation of cognitive function in aged mice with HDAC9 overexpression in forebrain glutamatergic neurons. (**A**) Schematic diagram of the HDAC9-Tg^flox-STOP-flox^ construct. (**B**) Genotyping results using different primers indicated in (A). (**C**) Breeding strategy to generate Emx1-ires-Cre;HDAC9-Tg^flox-STOP-flox^ (Emx1-ires-Cre;HDAC9^OE^) mice for glutamatergic neuron-specific HDAC9 overexpression. (**D**) HDAC9 mRNA expression levels in the hippocampus. Two-tailed unpaired t-test with Welch’s correction, *P* < 0.001. *n* = 6-9 mice per group. (**E**) HDAC9 and FLAG protein expression in the hippocampus. Two-tailed unpaired t-test, *t*_7_ = 11.29, *P* < 0.001. *n* = 4-5 mice per group. (**F**) Y-maze spontaneous alternation test at different ages. Young age - 6 months: spontaneous alternation rate, two-tailed unpaired t test, *t*_29_ = 0.466, *P* = 0.645; total arm entries, two-tailed unpaired t test, *t*_29_ = 1.407, *P* = 0.170. Middle age - 15 months: spontaneous alternation rate, two-tailed unpaired t test, *t*_28_ = 0.184, *P* = 0.856; total arm entries, Mann Whitney test, *P* = 0.009. Old age – 21 months: spontaneous alternation rate, two-tailed unpaired t test, *t*_17_ = 2.467, *P* = 0.025; total arm entries, Mann Whitney test, *P* = 0.017. *n* = 9-18 per group. (**G**) Novel object recognition (NOR) test. Middle age -15 months: exploration time of Emx1-ires-Cre;HDAC9^OE^ mice, two-tailed paired t test, *t*_12_ = 6.521, *P* < 0.001; exploration time of Emx1-ires-Cre littermate controls, Wilcoxon matched-pairs signed rank test, P < 0.001; discrimination index, Mann Whitney test, *P* = 0.363. Old age – 21 months: exploration time of Emx1-ires-Cre;HDAC9^OE^, two-tailed paired t test, *t*_8_ = 0.543, *P* = 0.602; exploration time of Emx1-ires-Cre littermate controls, Wilcoxon matched-pairs signed rank test, P = 0.221; discrimination index, two-tailed unpaired t test, *t*_17_ = 0.766, *P* = 0.454. *n* = 9-17 per group. (**H**) Social memory test. Left, schematic illustration of the social memory test setup. Middle-left, representative heat maps showing time spent in the test arena. Middle-right, quantification of exploration time with the familiar mouse (M1) versus the novel mouse (M2): Emx1-ires-Cre;HDAC9^OE^, two-tailed paired t test, *t*_7_ = 3.063, *P* = 0.0182. Emx1-ires-Cre, Wilcoxon matched-pairs signed rank test, *P* = 0.922. Right, social preference index. Mann Whitney test, *P* = 0.021. *n* = 8-10 per group. (**I**) mRNA expression levels of total *Bdnf*, *Bdnf* exon-IV and *Bdnf* exon-IV in the hippocampus. Total *Bdnf*, two-tailed unpaired t test, *t*_10_ = 3.193, *P* = 0.009; *Bdnf* exon-IV, two-tailed unpaired t test, *t*_10_ = 2.244, *P* = 0.049; *Bdnf* exon-IV, two-tailed unpaired t test, *t*_10_ = 1.298, *P* = 0.223. *n* = 5-7 per group. \**P* < 0.05, \*\**P* < 0.01, \*\*\**P* < 0.001. ns, no statistical significance.

To generate the knock-in line, the targeting vector was introduced into embryonic stem (ES) cells derived from the C57BL/6 strain. Correctly targeted ES clones were injected into albino C57BL/6J blastocysts, and the resulting chimeric founders were bred with C57BL/6J females to establish germline-transmitting heterozygous mice, hereafter referred to as HDAC9-Tg^flox-STOP-flox^ mice (Fig. 4B). To drive HDAC9 overexpression in the forebrain glutamatergic neurons, HDAC9-Tg^flox-STOP-flox^ mice were crossed with Emx1-ires-Cre knock-in mice (Fig. 4C), which have the endogenous *Emx1* locus directing expression of Cre recombinase predominantly to glutamatergic neurons in the hippocampus and neocortex ^88^. As expected, HDAC9 mRNA expression levels were increased in the hippocampus of Emx1-ires-Cre;HDAC9-Tg^flox-STOP-flox^ (hereafter Emx1-ires-Cre;HDAC9^OE^) mice compared to Emx1-ires-Cre control littermates (Fig. 4D). Elevated HDAC9 protein levels and FLAG-tag expression were further confirmed by western blotting (Fig. 4E).

To evaluate the impact of HDAC9 overexpression on age-related cognitive decline, we conducted a series of behavioral tests across different age groups. The Y-maze spontaneous alternation test, which can be used longitudinally to assess working memory in the same cohort of mice ^89–91^, was performed at 6, 15, and 21 months of age. In young adult (6-month-old) and middle-aged (15-month-old) mice, HDAC9 overexpression had no significant effect on short-term working memory, as indicated by similar alternation rates between Emx1-ires-Cre;HDAC9^OE^ and Emx1-ires-Cre mice (Fig. 4F). However, at 21 months of age, Emx1-ires-Cre control mice exhibited marked working memory deficits, with spontaneous alternation rates near chance level (50%). Age-matched Emx1-ires-Cre;HDAC9^OE^ mice showed higher alternation rates (Fig. 4F). Additionally, 15- and 21-month-old Emx1-ires-Cre;HDAC9^OE^ mice made more arm entries in the Y-maze than Emx1-ires-Cre littermates (Fig. 4F), suggesting increased exploratory activity. Short-term object recognition memory was assessed using the NOR test at 15 and 21 months of age. At 15 months, both Emx1-ires-Cre;HDAC9^OE^ mice and Emx1-ires-Cre littermates were able to discriminate between novel and familiar objects, with no significant differences in discrimination index (Fig. 4G). By 21 months, however, neither group displayed a preference for novel objects, and the discrimination index for both was near zero, indicating a loss of object recognition memory (Fig. 4G). Social recognition memory was further evaluated at 21 months of age. Emx1-ires-Cre mice spent similar amounts of time exploring familiar and novel conspecifics, indicating impaired social recognition memory. In contrast, Emx1-ires-Cre;HDAC9^OE^ mice spent significantly more time interacting with the novel mouse and demonstrated a higher social preference index (Fig. 4H), suggesting preserved social recognition memory. These findings indicate that HDAC9 overexpression in forebrain glutamatergic neurons can preserve both working and social memory in aged mice.

Given the critical role of BDNF in memory formation ^80–82^, we measured *Bdnf* mRNA levels in the hippocampus of Emx1-ires-Cre;HDAC9^OE^ mice compared with Emx1-ires-Cre littermates. We found that overexpression of HDAC9 in forebrain glutamatergic neurons increased the expression of total *Bdnf* mRNA. To determine whether specific *Bdnf* transcript variants were differentially affected, we next assessed the expression of exon-IV and exon-VI. HDAC9 overexpression significantly increased the levels of *Bdnf* exon-IV-containing transcripts, while *Bdnf* exon-VI transcript levels remained unchanged (Fig. 4I). These findings suggest that HDAC9 positively regulates *Bdnf* expression, particularly by modulating transcription of *Bdnf* exon-IV.

### HDAC9 expression is downregulated in the PFC and hippocampus in two mouse models of AD

To investigate whether HDAC9 contributes to the pathophysiology of AD, we first measured HDAC9 expression levels in the brain of 5-6-month-old 5xFAD mice. This AD model overexpresses human *APP* and *PSEN1* transgenes harboring five familial AD-linked mutations, specifically in neurons. HDAC9 mRNA levels were decreased in both the PFC and hippocampus of 5xFAD mice compared to WT littermate controls (Fig. 5A). Western blot analysis confirmed a corresponding reduction in HDAC9 protein expression in the PFC of 5xFAD mice (Fig. 5A). We also examined HDAC9 mRNA expression in the hippocampus of APP/PS1 mice, another widely used AD model. These mice express a chimeric mouse/human *APP* carrying Swedish mutations (K595N/M596L) and a mutant human *PSEN1* (ΔE9) under the control of mouse prion protein promoter elements ^92,93^. Similar to 5xFAD mice, HDAC9 mRNA expression was decreased in the hippocampus of APP/PS1 mice compared to WT littermate controls; a decreasing trend was also observed in the PFC of APP/PS1 mice (Fig. 5B). Collectively, these findings indicate that HDAC9 expression is downregulated in the brain of two AD mouse models, suggesting a potential role for HDAC9 in AD-related phenotypes.

**Figure 5.**
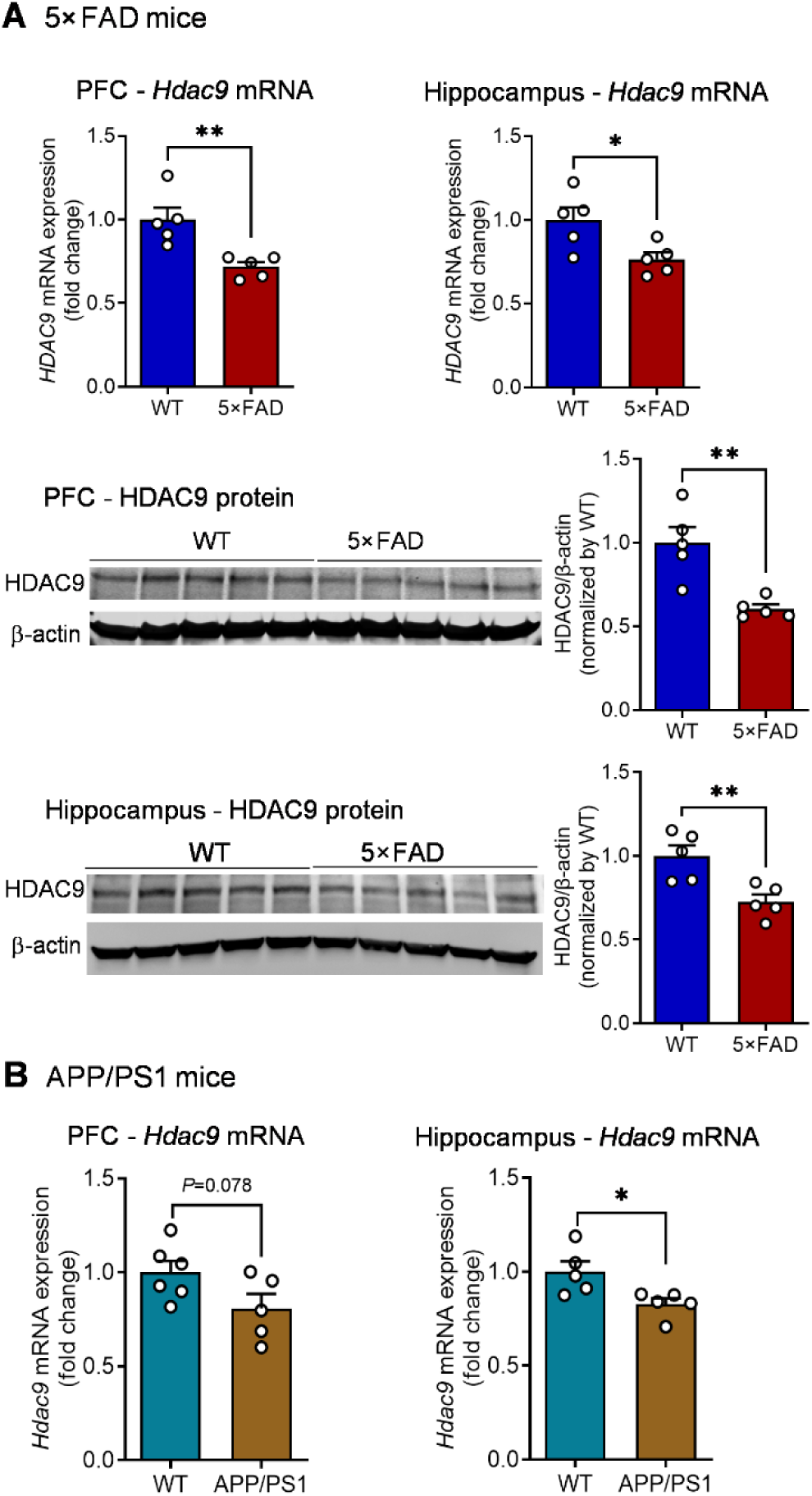
HDAC9 expression in the brain of mouse models of AD. (**A**) 5xFAD mice. HDAC9 mRNA and protein expression levels in the prefrontal cortex (PFC) and hippocampus of 5xFAD mice and WT littermate controls. Upper, mRNA expression of HDAC9 in the PFC (two-tailed unpaired t test, *t*_8_ = 3.687, *P* = 0.006) and hippocampus (two-tailed unpaired test, *t*_8_ = 2.707, *P* = 0.027). Middle, Western blot showing HDAC9 protein expression levels in the PFC (two-tailed unpaired t test, *t*_8_ = 4.087, *P* = 0.004). Bottom, Western blot showing HDAC9 protein levels in the hippocampus (two-tailed unpaired t test, *t*_8_ = 3.545, *P* = 0.008). *n* = 5 mice per group. (**B**) HDAC9 mRNA expression levels in the PFC and hippocampus of APP/PS1 mice and WT littermate controls. Left, PFC (two-tailed unpaired t test, *t*_9_ = 1.992, *P* = 0.078). Right, hippocampus (two-tailed unpaired t test, *t*_8_ = 2.706, *P* = 0.027). *n* = 5-6 mice per group. \**P* < 0.05, \*\**P* < 0.01.

### Neuronal HDAC9 overexpression mitigates cognitive and synaptic deficits in 5xFAD mice

To assess whether overexpression of HDAC9 could ameliorate cognitive and synaptic deficits in 5xFAD mice, we crossed HDAC9-Tg^flox-STOP-flox^ mice with 5xFAD mice to generate 5xFAD/ HDAC9-Tg^flox-STOP-flox^, 5xFAD and WT offspring (Fig. 6A). Cre recombinase was delivered via tail-vein injection using AAV-PHP.eB, a serotype capable of crossing the blood–brain barrier for transduction of the brain ^94^. Cre was fused to GFP and driven by the human synapsin promoter, enabling neuron-specific expression and visualization. An AAV-PHP.eB-GFP vector lacking Cre recombinase was used as a control. At 5–6 months of age, 5xFAD/HDAC9-Tg^flox-STOP-flox^ mice received AAV-PHP.eB-GFP-Cre injection (hereafter referred to as 5xFAD/HDAC9^OE^). Age-matched 5xFAD and a subset of 5xFAD/HDAC9-Tg^flox-STOP-flox^ mice received the control AAV-PHP.eB-GFP injection (hereafter referred to as 5xFAD), as did WT control mice (Fig. 6B). Widespread expression of GFP-Cre was observed in the brain, including the hippocampus and cortex (Fig. 6C). Western blot analysis confirmed Cre-induced expression of FLAG-tagged HDAC9 and increased HDAC9 protein levels in the hippocampus compared with control AAV-injected mice (Fig. 6D). Four weeks after AAV injection, mice underwent a battery of behavioral tests (Fig. 6B). In the Y-maze test, 5xFAD mice exhibited a lower spontaneous alteration rate compared to WT mice, indicative of impaired spatial working memory (Fig. 6E). HDAC9 overexpression induced by AAV-PHP.eB-GFP-Cre improved spatial working memory, as evidenced by an increased alternation rate in the Y-maze. Additionally, HDAC9 overexpression reduced the total arm entries in 5xFAD mice, indicating decreased hyperactivity (Fig. 6E). Object recognition memory was evaluated following a training session with a 2 h retention interval. 5xFAD mice failed to distinguish between familiar and novel objects, exhibiting a lower discrimination index compared to WT mice. In contrast, HDAC9 overexpression rescued this deficit, normalizing the discrimination index (Fig. 6F). Reduced locomotor activity in 5xFAD mice by HDAC9 overexpression was also observed in the open field, as indicated by a decreased total distance travelled (Fig. 6G). Furthermore, anxiety-like behavior was evaluated using the elevated plus maze test. 5xFAD mice spent more time in the open arms compared to WT mice. This abnormal behavior was normalized by HDAC9 overexpression (Fig. 6H).

**Figure 6.**
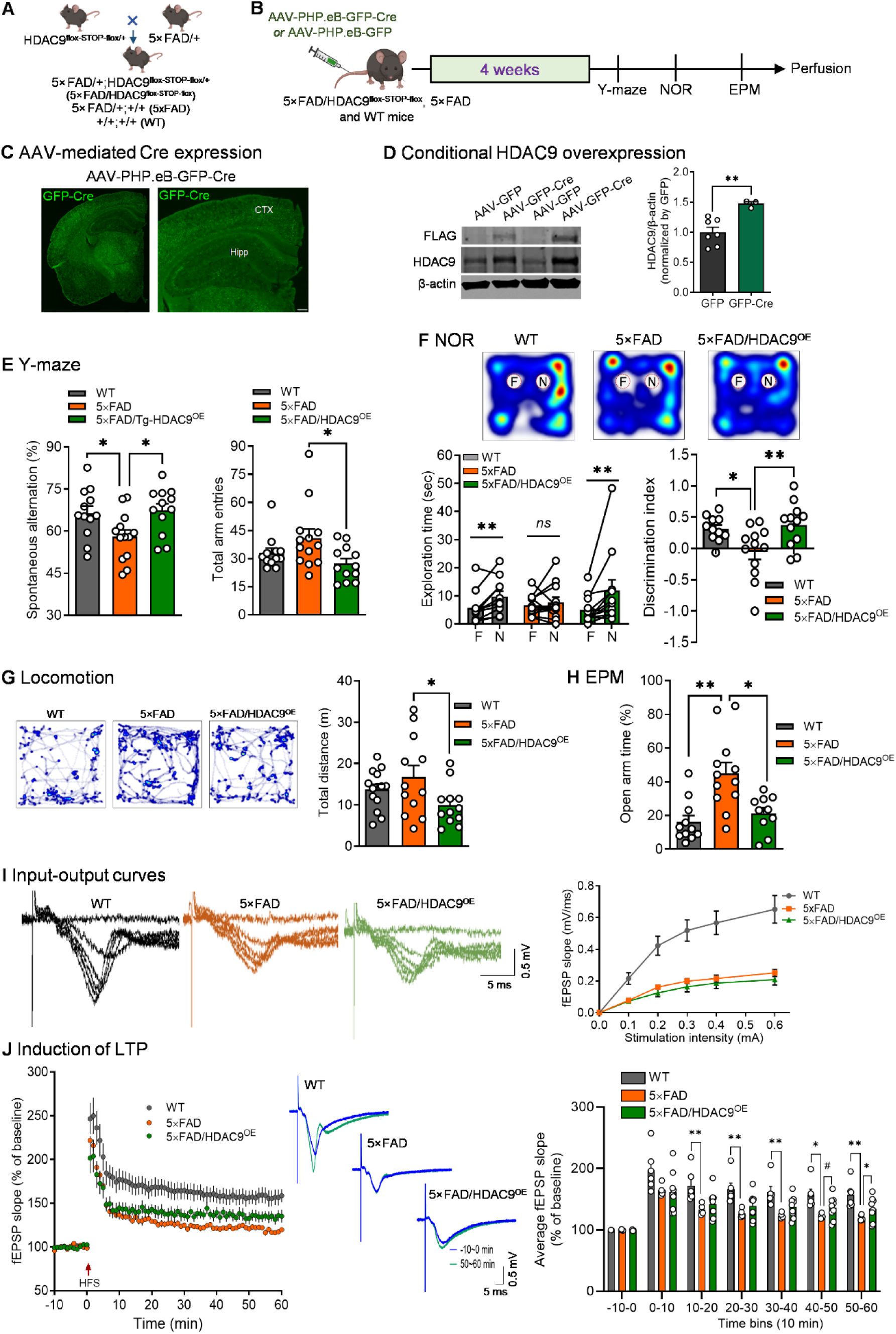
Reversal of behavioral and synaptic impairments of 5xFAD mice by neuronal HDAC9 overexpression. (**A**) Breeding strategy to generate 5xFAD/HADC9-Tg^flox-STOP-flox^ mice. (**B**) Schematic illustration of intravenous AAV injection and experimental timeline. (**C**) Representative image showing GFP-Cre expression in the mouse brain. Scale bar, 200 µm. (**D**) AAV-PHP.eB-mediated HDAC9 overexpression and FLAG expression in the hippocampus. Two-tailed unpaired t test, *t*_8_ = 3.575, *P* = 0.007. (**E**) Y-maze test. Left, spontaneous alternation rate, one-way ANOVA, *F*_(2, 34)_ = 4.473, *P* = 0.019; right, total arm entries, Kruskal-Wallis test, *P* = 0.043. *n* = 12-13 mice per group. (**F**) Novel object recognition (NOR) test. Exploration time of WT mice: Wilcoxon matched-pairs signed rank test, *P* = 0.0098. Exploration time of 5xFAD mice: two-tailed paired t-test, *t*_12_ = 0.648, *P* = 0.531. Exploration time of 5xFAD/HDAC9^OE^ mice: Wilcoxon matched-pairs signed rank test, *P* = 0.005. Discrimination index: One-way ANOVA, *F*_(2, 32)_ = 5.549, *P* = 0.009. *n* = 11-12 mice per group. (**G**) Locomotor activity. One-way ANOVA, *F*_(2, 35)_ = 3.068, *P* = 0.059. *n* = 12-14 mice per group. (**H**) Elevated plus maze (EPM) test. Percentage of time spent in the open arms during the test. Kruskal-Wallis test, *P* = 0.002. *n* = 10-12 mice per group. (**I**) Input–output curves of fEPSPs. Left, representative traces of fEPSPs recorded in the CA1 region in response to increasing stimulation intensities delivered to Schaffer collateralls. Right, quantitative Input-output curves showing the relationship between stimulation intensity and fEPSP slope. Two-way repeated measures ANOVA: treatment, *F*_(2, 18)_ = 16.05, *P* < 0.001; stimulation, *F*_(5, 90)_ = 98.85, *P* < 0.001; genotype × treatment, *F*_(10, 90)_ = 15.57, *P* < 0.001. (**J**) Long-term potentiation (LTP) recordings at CA3-CA1 synapses. Left, time course of fEPSPs slopes recorded before and after high-frequency stimulation (HFS) of Schaffer collaterals. Two-way repeated measures ANOVA: treatment, *F*_(2, 17)_ = 5.600, *P* = 0.014; time, *F*_(70, 1190)_ = 82.81, *P* < 0.001; treatment × time, *F*_(140, 1190)_ = 3.062, *P* < 0.001. Middle, representative traces recorded 10 min before HFS (baseline) and 50 min after HFS. Right, average fEPSP slope calculated in 10-min time bins before and after LTP induction. -10-0 min, One-way ANOVA, *F*_(2, 17)_ = 0.720, *P* = 0.501; 0-10 min, One-way ANOVA, *F*_(2, 17)_ = 3.127, *P* = 0.070; 10-20 min, One-way ANOVA, *F*_(2, 17)_ = 5.555, *P* = 0.014; 20-30 min, Kruskal-Wallis test, *P* = 0.008; 30-40 min, Kruskal-Wallis test, *P* = 0.003; 40-50 min, Brown-Forsythe ANOVA test, *P* = 0.013; 50-60 min, Brown-Forsythe ANOVA test, *P* = 0.008. WT: *n* = 6 slices of 4 mice; 5xFAD: *n* = 5 slices of 4 mice; 5xFAD/HDAC9^OE^: *n* = 9 slices of 7 mice. \**P* < 0.05, \*\**P* < 0.01, ^#^*P* = 0.053.

To determine whether the improved cognitive function resulting from HDAC9 overexpression is associated with improved synaptic plasticity, a separate cohort of 5xFAD and 5xFAD/HDAC9-Tg^flox-STOP-flox^ mice received tail vein injections of either AAV-PHP.eB-GFP-Cre or the control AAV-PHP.eB-GFP. Four weeks post-injection, acute hippocampal slices were prepared for recordings of fEPSPs in the CA1 region in response to stimulation of the Schaffer collateral pathway. Compared to WT controls, 5xFAD mice exhibited reduced fEPSP responses across a range of stimulus intensities, indicating impaired basal synaptic transmission. HDAC9 overexpression in 5xFAD mice, achieved via AAV-PHP.eB-GFP-Cre, didn’t significantly alter input-output responses compared to 5xFAD mice receiving the AAV-PHP.eB-GFP. LTP was then induced using high-frequency stimulation to assess synaptic plasticity. As expected, 5xFAD mice showed impaired LTP compared to WT controls. Notably, HDAC9 overexpression partially rescued this deficit, as 5xFAD/HDAC9-overexpressing mice exhibited significantly improved LTP maintenance relative to 5xFAD controls (Fig. 6J), suggesting that HDAC9 enhances synaptic plasticity in this AD model.

### Neuronal overexpression of HDAC9 reduces Aβ plaque accumulation in 5xFAD mice

To determine whether HDAC9 overexpression influences neuropathology of 5xFAD mice, brains were harvested following behavioral testing and processed for immunohistochemical analysis. Aβ deposition was assessed using the 6E10 antibody, which recognizes the N-terminal amino acids 1– 16 of APP and Aβ peptides, thus detecting multiple forms of Aβ. Quantitative analysis of Aβ plaque deposition was performed in three brain regions involved in regulating learning and memory and are affected by AD ^95–97^, including the mPFC, hippocampus and entorhinal cortex (EC). Within the mPFC, the prelimbic (PL) and infralimbic (IL) subregions have similar cellular composition and distinct functions in cognitive processes ^98^. HDAC9 overexpression reduced Aβ plaque load in both the PL and IL subregions of mPFC compared to 5xFAD controls (Fig. 7A). In the hippocampus, the Aβ plaque load was quantified in the CA1, CA3 and dentate gyrus (DG). HDAC9 overexpression led to significant reductions in plaque load across all subregions, with the greatest decrease observed in CA3 (CA1: 44.34%, CA3: 62.92%, DG: 58.91%) (Fig. 7B). Aβ plaque accumulation was also markedly decreased in the EC of 5xFAD/HDAC9^OE^ mice (Fig. 7C). Collectively, these findings suggest that neuronal HDAC9 overexpression mitigates Aβ pathology across multiple brain regions in 5xFAD mice.

**Figure 7.**
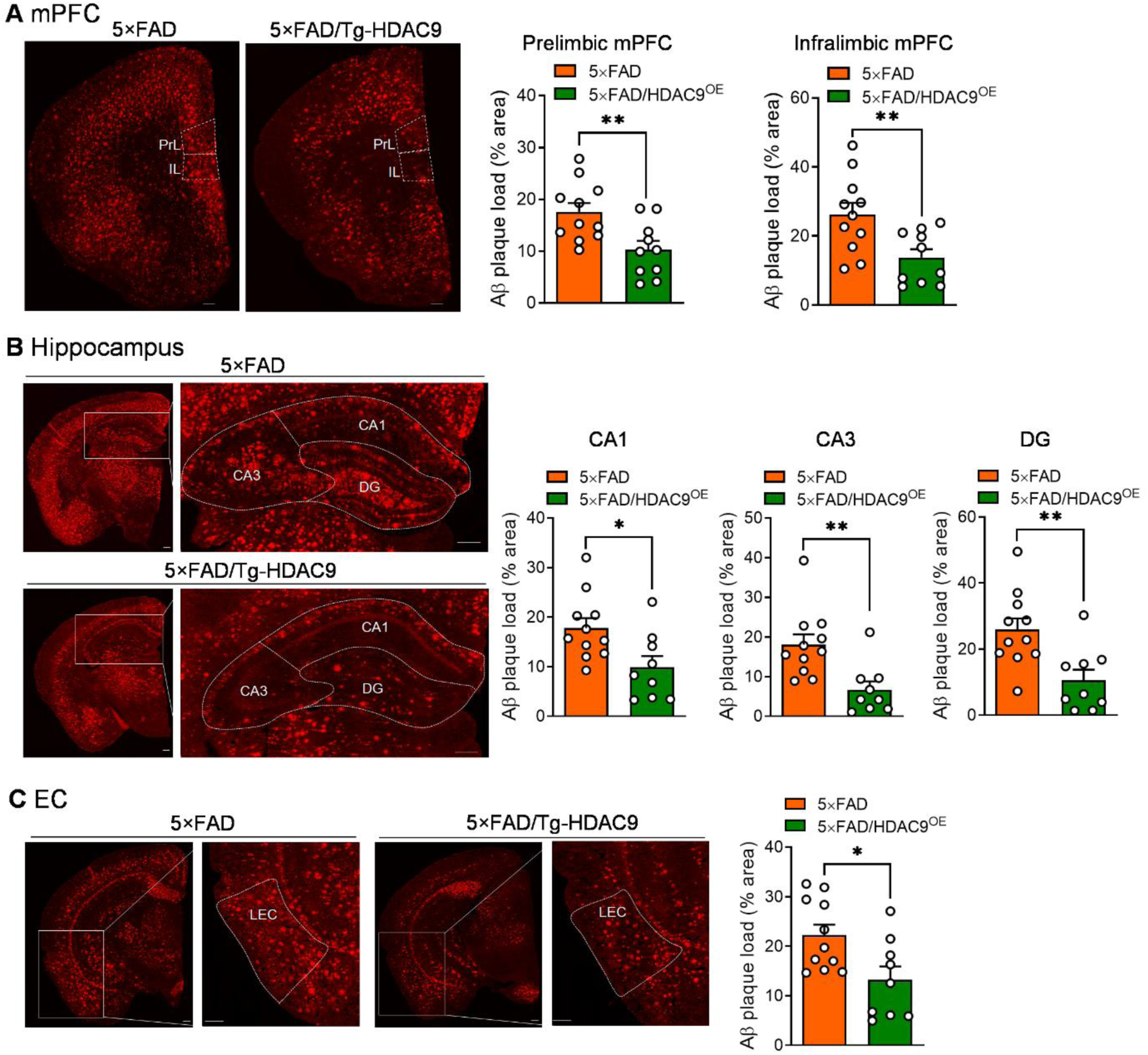
Reduction of Aβ deposition by neuronal overexpression of HDAC9 in 5xFAD mice. Representative images (left) and quantitative analysis (right) of Aβ plaque deposition in the medial prefrontal cortex (mPFC, **A**), hippocampus (**B**) and entorhinal cortex (EC, **C**). Prelimbic mPFC: two-tailed unpaired t test, *t*_19_ = 3.015, *P* = 0.007. Infralimbic mPFC: two-tailed unpaired t test, *t*_19_ = 2.925, *P* = 0.009. CA1: two-tailed unpaired t test, *t*_18_ = 2.631, *P* = 0.017. CA3: Mann Whitney test, P = 0.003. Dentate gyrus (DG): two-tailed unpaired t test, *t*_18_ = 3.190, *P* = 0.005. EC: two-tailed unpaired t test, *t*_18_ = 2.647, *P* = 0.016. *n* = 9-11 mice per group. \**P* < 0.05, \*\**P* < 0.01.

## DISCUSSION

Over the course of aging, cognitive abilities decline as neuropathological changes build up, particularly in brain regions such as the hippocampus and prefrontal cortex, which are vulnerable to age-related degeneration ^72–74^. Impaired synaptic plasticity is a characteristic neurobiological change during aging ^78,99^ and strongly correlates with cognitive decline ^3,100,101^. While many studies have shown that pharmacological inhibition of HDACs improves cognitive function and synaptic plasticity ^22–30,102^, the limited isoform selectivity of currently available HDAC inhibitors remains a major obstacle to defining the distinct roles of individual HDAC isoforms ^19,35^. Moreover, the diverse and cell-specific functions of HDAC isoforms remain largely unexplored. Therefore, investigating endogenous HDAC expression and regulation within specific cell types during aging and AD is essential to elucidate their distinct roles and influence on cognitive processes in physiological and pathological states.

In this study, we characterized HDAC9 expression in neurons, microglia and astrocytes, demonstrating that HDAC9 is exclusively expressed in neurons in both human and mouse brains, with no detectable expression in microglia or astrocytes. These findings suggest that HDAC9 functions as a cell type–specific epigenetic or transcriptional regulator restricted to neurons. Additionally, we assessed HDAC9 expression at various ages and found that it decreases with age in the hippocampus and PFC, regions critical for learning and memory, suggesting that altered HDAC9 expression may play a role in age-related neuronal dysfunction and cognitive decline. A reduction in HDAC9 expression at both the mRNA and protein levels was also observed in the hippocampus and PFC of 5xFAD mice, a widely used model of AD ^71^. A similar reduction at the mRNA level was confirmed in the same brain regions of APP/PS1 mice, another commonly used AD model. These findings are consistent with reports from human AD studies ^53,55,56^. In contrast, a recent study reported elevated HDAC9 protein expression in the forebrain of APP/PS1 mice, although the specific brain regions analyzed and details of the mouse line were not provided ^103^. This discrepancy could be due to the specificity of HDAC9 antibodies used and the brain regions examined. Collectively, these findings indicate that HDAC9 is dynamically regulated during aging and in the context of AD pathology. Moreover, our results from HDAC9 knockout mice, which exhibited deficits in cognitive performance and synaptic plasticity, support a causal relationship between HDAC9 downregulation and both cognitive and synaptic impairments. Specifically, loss of HDAC9 led to impaired spatial working memory and object recognition memory. Given the critical role of CA1 neurons in the formation, consolidation, and retrieval of hippocampal-dependent memories ^104,105^, we further investigated the region-specific role of HDAC9 in adult CA1 neurons using AAV-Cre-mediated gene knockdown. We found that mice with CA1-specific HDAC9 knockdown exhibited significant cognitive deficits, reinforcing the notion that HDAC9 is required for hippocampal function and memory processing. Given that synaptic plasticity is a fundamental mechanism underlying learning and memory ^106–108^, we further assessed the impact of loss of HDAC9 on synaptic function. HDAC9 knockout mice failed to maintain hippocampal LTP, indicating impaired synaptic plasticity. The concurrent reduction in synaptic plasticity and cognitive performance further supports that HDAC9 is required for maintaining synaptic integrity, plasticity and cognitive function. Its loss compromises both the functional and structural substrates of memory, likely contributing to age-related cognitive decline and the synaptic and cognitive deficits associated with AD.

These observations raise the critical question of whether increasing HDAC9 expression can preserve cognitive function during aging. Glutamatergic neurons in the cortex and hippocampus are particularly vulnerable to age-related changes ^99,109–111^, and their structural and functional alterations are closely associated with cognitive decline in aging ^99,112–114^. To investigate the role of HDAC9 in this neuronal population in the context of age-related cognitive decline, we generated a conditional knock-in mouse line that overexpresses HDAC9 specifically in forebrain glutamatergic neurons and characterized their behavioral phenotypes from young adulthood through middle age to old age. Our results showed that HDAC9 overexpression did not significantly affect cognitive performance at 6 or 15 months of age, indicating that elevating HDAC9 levels in glutamatergic neurons do not enhance cognition in young or middle-aged mice. However, by 21 months of age, control mice exhibited prominent cognitive deficits across multiple cognitive domains, including impairments in spatial working memory, novel object recognition, and social recognition memory. Remarkably, HDAC9 overexpression preserved spatial working memory and improved social recognition memory in aged mice, suggesting a protective effect against age-related decline. However, HDAC9 overexpression did not ameliorate the deficits observed in novel object recognition, implying that its protective effects may be task-specific. These findings suggest that modulating HDAC9 expression could be a promising strategy to enhance cognitive resilience in the aging brain.

The observation of HDAC9 downregulation in the hippocampus and PFC of AD mouse models led to the investigation of whether restoring HDAC9 expression in neurons could rescue or mitigate the behavioral and neuropathological phenotypes associated with AD. To address this question, we utilized the blood-brain barrier–penetrant AAV-PHP.eB vector ^94^ to deliver Cre recombinase driven by a neuron-specific promoter, thereby inducing HDAC9 overexpression in neurons of 5xFAD mice. Our findings demonstrated that neuron-specific HDAC9 overexpression was sufficient to rescue the cognitive deficits observed in 5xFAD mice. Furthermore, the impaired synaptic plasticity in 5xFAD mice was partially reversed by HDAC9 overexpression, pointing to a functional rescue of synaptic plasticity. Strikingly, we also observed a reduction in Aβ plaque burden in multiple brain regions critically involved in memory and highly vulnerable in AD ^115–120^. This reduction in Aβ pathology suggests that HDAC9 may exert a disease-modifying effect, either by directly influencing Aβ production/clearance or indirectly through improved neuronal function. The mechanisms by which HDAC9 regulates cognitive and synaptic function likely involve neurotrophic signaling pathways. In HDAC9 knockout mice, impairments in cognitive and synaptic function and structure were accompanied by a decrease in BDNF expression in the hippocampus. Conversely, HDAC9 overexpression in forebrain glutamatergic neurons increases BDNF levels in the hippocampus. BDNF is a key modulator of synaptic plasticity and memory formation, and its dysregulation has been implicated in both aging-related cognitive decline and AD ^80–82^. The *BDNF* gene in mice consists of nine distinct exons and nine individual functional promoters ^121,122^. While all transcripts encode an identical BDNF protein, exon-specific transcription is controlled by distinct promoters, allowing for precise spatial and temporal regulation of BDNF expression in response to specific stimuli ^83^. Each BDNF isoform is associated with a distinct set of functional outcomes ^83^. Notably, BDNF exon-IV and exon-VI transcripts are driven by neuronal activity ^84,85^, and dysregulation of these exons has been linked to memory disorders ^80,86^. Our analysis of exon-IV and exon-VI expression revealed that loss of HDAC9 decreases exon-IV expression without affecting exon-VI expression. This suggests that reduced exon-IV transcription may cause a reduction in activity-dependent BDNF expression in the hippocampus, contributing to impaired cognitive function ^80,86^. In contrast, HDAC9 overexpression in glutamatergic neurons of aged mice may improve synaptic plasticity and cognitive function by increasing BDNF expression. In the context of AD, a marked reduction in BDNF levels in the cortex and hippocampus has been reported ^123–125^. This decrease in BDNF is believed to disrupt synaptic plasticity and contribute to cognitive decline in AD mouse models. In addition to its neurotrophic effects, BDNF has been shown to reduce Aβ levels, potentially through a mechanism involving increased α-secretase activity ^126^. Given these roles, we speculate that BDNF may mediate, at least in part, the HDAC9-induced reduction in Aβ accumulation observed in 5xFAD mice. Further studies will be needed to determine whether HDAC9 overexpression enhances BDNF signaling and whether this contributes to the observed cognitive and neuropathological improvements.

These findings support a new concept that HDAC9 downregulation may represent a mechanistic link between aging-related cognitive decline, synaptic vulnerability, and the progression of AD. While pharmacological inhibition of HDACs has generally been associated with enhanced cognitive function ^6,127^, our results reveal a distinct, isoform-specific role for HDAC9: neuronal overexpression of HDAC9 preserves cognitive function during aging and mitigates AD-associated cognitive deficits and neuropathology. Future studies are warranted to elucidate the downstream molecular pathways regulated by HDAC9. Together with the observed HDAC9 downregulation in aging and AD models, our data suggest that HDAC9 is not only necessary for maintaining cognitive and synaptic function but also sufficient to preserve synaptic cognitive performance in the aging and AD brain. These findings position HDAC9 as a promising therapeutic target for mitigating age-related cognitive decline and AD.

## ACKNOWLEDGMENT

This project was supported by the National Institute of Health (NIH) grants (AG076235, AG064895, AG083841 and AG062166). We gratefully acknowledge the Augusta University Genome Editing Core for generating the knock-in mouse line and the Neuropathology Core at the Emory Alzheimer’s Disease Research Center for providing human post-mortem brain tissues.

## DATA AVAILABILITY STATEMENT

The datasets generated during the current study are available from the corresponding authors upon request.

